# Effects of a novel thermophilic cellulose-degrading agent on the quality of compost and change in microbial community of garden waste

**DOI:** 10.1101/688853

**Authors:** Zhouzhou Fan, Zhenzhen Jia, Yongshuang Li, Peng Lian, Xiawei Peng

## Abstract

Knowledge about the microbial communities in composting has advanced, but definitive knowledge concerning the application of actinomycetal communities in garden waste composting is still lacking. In this study, we compared the effects of amending compost with mixed agent M1 (five high-degradability strains) and other agents on the physicochemical indices and microbial community succession. The results showed that Pile A (only applying M1), exhibited a pH closer to neutral, the complete degradation of organic matter, and the highest remaining levels of nitrogen, phosphorus, and potassium. The seed germination rate, root length, and seed germination index values were significantly higher in piles amended with M1 and/or commercially available agents than in piles without exogenous microbial agents. Analyzing the microbial communities, these treatments were dominated by Proteobacteria, Firmicutes, Actinobacteria, and Bacteroidetes during composting. The amount of *Streptomyces* was negatively correlated with the carbon/nitrogen ratio and positively correlated with total phosphorus and total potassium. Adding M1 increased microbial diversity, and the dominant microbial communities at the end of composting were similar to those found in the commercial microbial inoculum. Overall, agent M1 can shorten the composting process and increase the extent of degradation. This research provides additional insights into the potential function of Actinomycetes in compost ecology.

## Introduction

Waste management is one of the main issues affecting all developing nations [1]. In China, with the acceleration of urbanization, the amount of green area coverage is increasing, but tons of organic solid waste are produced with no means for effective decomposition, representing added stress to the environment [2]. The main components of garden waste, lignin and cellulose, are recalcitrant to degradation, leaving only landfilling and direct burning as traditional disposal methods. Traditional landfilling and incineration methods not only use land resources, but also cause other problems, such as dust particle production, soil and groundwater contamination, pathogenic bacteria growth, and air pollution [3]. To lessen the effects of garden waste disposal on the environment and mitigate environmental pollution, biological composting employs microbial action in waste piles to transform organic solid waste into stable humus and organic fertilizer. Biological composting can directly transform large amounts of organic waste for compost production; however, the unreliability of the quantity and biodegradability of the indigenous functional microbial community in compost often leads to low composting efficiency and undesirable compost quality [4].

Many studies have shown that inoculation with exogenous microorganisms is an effective method for the biodegradation of organic matter and accelerates the composting process [5-6]. Through contrast tests, Gong [7] studied the effects of thermophilic bacteria on sludge composting, which increased rate of temperature rise and effectively enhanced the degradation of organic matter. Meanwhile, Bonito *et al* [8] reported that fungi were associated with composting of organic municipal waste. In addition, Amira *et al* [9] showed that the application of fungi to compost resulted in higher xylanase and cellulase activity, consequently leading to the rapid degradation of cellulose and hemicellulose. However, many studies have not assessed the amendment of compost with actinomycetal communities, which are abundant in environments rich in lignocellulose and have essential roles in nutrient recycling. During composting, Actinomycete amendment might alter microbial communities to subsequently alter lignocellulose degradation. Wang *et al.* [10] found that Actinomycetes account for 18–86% of the bacterial community, which peaked during the compost maturation phase. Furthermore, exogenously added agents degrade cellulose through the synergistic actions of a series of enzymes, promoting the degradation of organic matter, shortening the composting cycle, and accelerating compost maturation [11-13]. High-throughput sequencing is the most effective technique for detecting microbial diversity, and has been widely applied in composting analyses.

Understanding such microbial interactions and dynamics could reveal the mechanisms driving material transformation and the overall process of compost maturation. Most studies have concluded that the effect of the addition of a single strain to compost is lower than that of mixed strains, because the degradation of organic matter requires a combination of many enzymes secreted by different microbes [14-15]. Nevertheless, few studies have added different microbial agents to individual compost piles, especially carefully screened agents, to compare the effects with natural compost on microbial communities. Therefore, in this study, we isolated Actinomycetes species from compost comprising garden waste and chicken manure under high temperatures with efficient, thermotolerant cellulase production to create a mixed culture, M1 (containing *Streptomyces thermoviolaceus*, *S. thermodiastaticus*, *S. thermocarboxydus*, *S. albidoflavus*, and *S. thermovulgaris*). Then, we assessed the influence of M1 on the quality of compost and change in microbial community during the composting process. The results will guide the rapid decomposition of organic solid waste and provide important information for optimizing inoculation technology and improving fermentation processes.

## Materials and Methods

### Composting materials

Five thermophilic cellulose-degrading strains with high enzyme productivity were isolated from compost from garden waste and chicken manure under high temperature, using Congo red for the initial screening and enzyme-producing ability assessments for additional screenings. The bacterial strains *S. thermoviolaceus*, *S. thermodiastaticus*, *S. thermocarboxydus*, *S. albidoflavus*, and *S. thermovulgaris* were identified by 16S rRNA gene sequence identification. The five strains have been deposited in the Chinese General Microbiological Culture Collection Center (No. 12133–12137) and their 16S rRNA gene sequences have been deposited into the NCBI GenBank database (accession numbers MF114295–MF114399). The strains showed no antagonism with each other; therefore, the strains were diluted during the logarithmic growth phase to an optical density at 600 nm of 0.2, and were then mixed in equal volumes for fermentation. The specific production of M1 agent is described in detail by Feng *et al* [5] A common commercial agent (Crude fiber degrading agent produced by the company of Wei Yuan Biology) was chosen for comparison. The viable cell numbers of agent M1 and the commercial agent was 10 ^9^ /g. A mixture of fresh sheep manure and garden green waste at a weight ratio of 3:1 was used as the compost and was supplied by a local farmer in Changping District, Beijing, China. Four parallel piles, each containing approximately 2000 kg of dry compost, were formed as the control (CK) and treatment (A, B, and C) groups. The treatments for each pile are listed in Table 1. To reduce the effects of non-experimental factors, all conditions were kept constant except the added microbial agent.

**Table 1.**
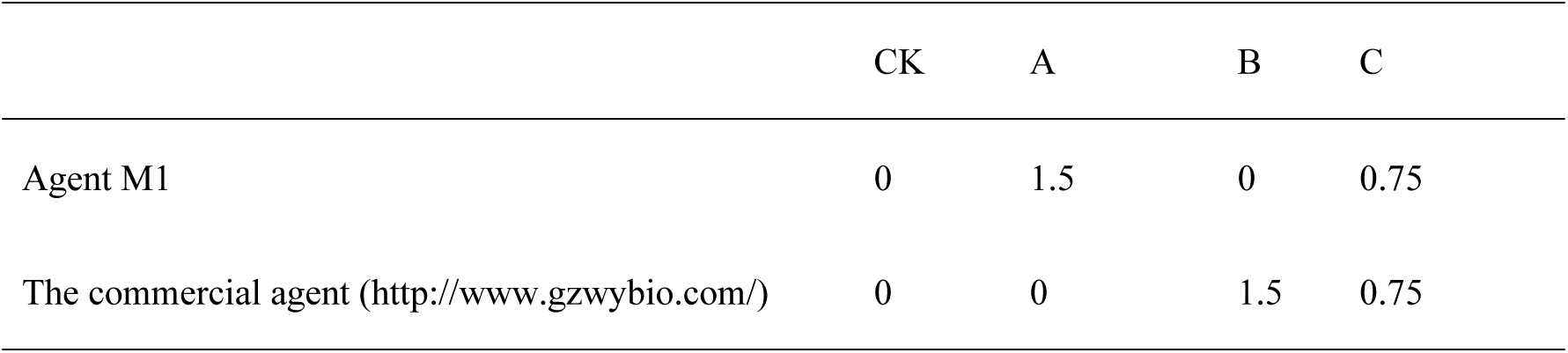
Microbial agent amendment in the four composting groups (unit: kg).

### Composting process and collection

The experiment was performed outdoors at the Aobei Merrill Biological Fertilizer Research Center (Beijing, China). The initial water content was determined by oven drying representative samples of each group, and was adjusted to the range of 55–60%. The compost was placed in piles (length: 3 m, width: 1.5 m, height: 1 m). The piles first were turned manually three times every 3 days, and turned three times every 7 days thereafter for a total of 40 days. Samples were taken at days 3, 6, 9, 12, 15, 18, 21, 25, 30, 35, and 40 during composting. At each sampling time, approximately 200 g of material was taken from three points in layer 1 (∼20 cm), layer 2 (∼50 cm), and layer 3 (∼80 cm) and thoroughly mixed into one sample. Three parallel samples were taken from each pile. In addition, 100 g subsamples collected on day 3 (mesophilic period), day 6 (early thermophilic period), day 12 (middle thermophilic period), day 18 (late thermophilic period), and day 40 (cooling period) were labeled as samples 1, 2, 3, 4, and 5, respectively, and stored at −80°C until DNA extraction for the microbial diversity analysis. The remainder of the material was used for the physiochemical analysis.

### Physicochemical analysis

Before the piles were turned manually, the temperature of the final samples and environment was measured with a digital thermometer at 10:00 during the composting process. The measurement position is in the middle of the composting. The other components of the compost were determined by referring to the routine methods described by Bao [16]. pH was measured in an aqueous suspension of the final sample (1:10, w/v, sample/water ratio) using a pH meter. The organic carbon and total nitrogen (TN) contents were measured via the dichromate oxidation and Kjeldahl methods, respectively. The total phosphorus (TP) content was analyzed colorimetrically following the ammonium molybdate method. Total potassium (TK) was determined using the flame photometer method. The germination assay was assessed with the seed germination rate (SGR), root length (RL), and seed germination index (GI) of the cabbage, as described previously [17]. GI was calculated as follows: GI (%) = (mean number of germinated seeds in the treatment × mean RL in the treatment × 100%) / (mean number of germinated seeds in water × mean RL in water).

### Microbial diversity analysis

DNA was extracted from compost samples using the FastDNA® Spin Kit for Soil (MP Biomedicals, Illkirch, France) following the manufacturer’s instructions. The concentration and purity of the extracted DNA were examined with a Nanodrop® ND-1000 spectrophotometer and agarose gel electrophoresis. The 16S rRNA genes of V4 hypervariable regions in the qualified samples were amplified using the special primer set 515F (5′-GTGCCAGCMGCCGCGG-3′) and 907R (5′-CCGTCAATTCMTTTRAGTTT-3′). PCR products were quantified and mixed in equal concentrations with a TBS-380 fluorescence meter (Turner Biosystems) and sequenced on an MiSeq PE300 (Illumina, San Diego, CA, USA). All raw Illumina MiSeq sequencing data is deposited in the NCBI Sequence Read Archive database (accession number SRP113350). The paired-end sequences were optimized using Trimmomatic and FLASH and classified into operational taxonomic units (OTUs) according to biological information at a 97% identity threshold, so that almost all OTUs could represent the true sequences. Rarefaction curves and Shannon and Chao indices were determined by RDP classifier (confidence threshold = 0.7) in QIIME [18]. Fast UniFrac was used to integrate related samples into single analyses. According to the bray-curtis method to perfom NMDS analysis using Vegan [19].

## Results

### The changes in chemical and biological indicators of composting maturity

#### Temperature profiles

Fig. 1(a) shows the temperature changes during the composting of piles A, B, C, and CK. The ambient temperature variation throughout the composting period was between 17°C and 26°C. The evolution of temperature in the compost piles was divided into three periods: mesophilic period, thermophilic period, and cooling period. An increase in compost pile temperature was observed soon after composting started, and the high temperature stage (> 50°C) of groups A, B, C, and CK began on days 3, 4, 5, and 6, respectively. In pile A, a fast increase in temperature was observed, reaching a maximum temperature of 72°C, and the thermophilic phase (> 60°C) lasted for 14 days, after which the temperature decreased rapidly to reach the cooling period (day 24). In piles B and C, gradual increases in temperature were observed; they reached the thermophilic phase with maximum temperatures of 66°C and 64°C, respectively, on day 18. CK exhibited a significantly lower maximum temperature and thermophilic phase (> 50°C).

**Fig 1.**
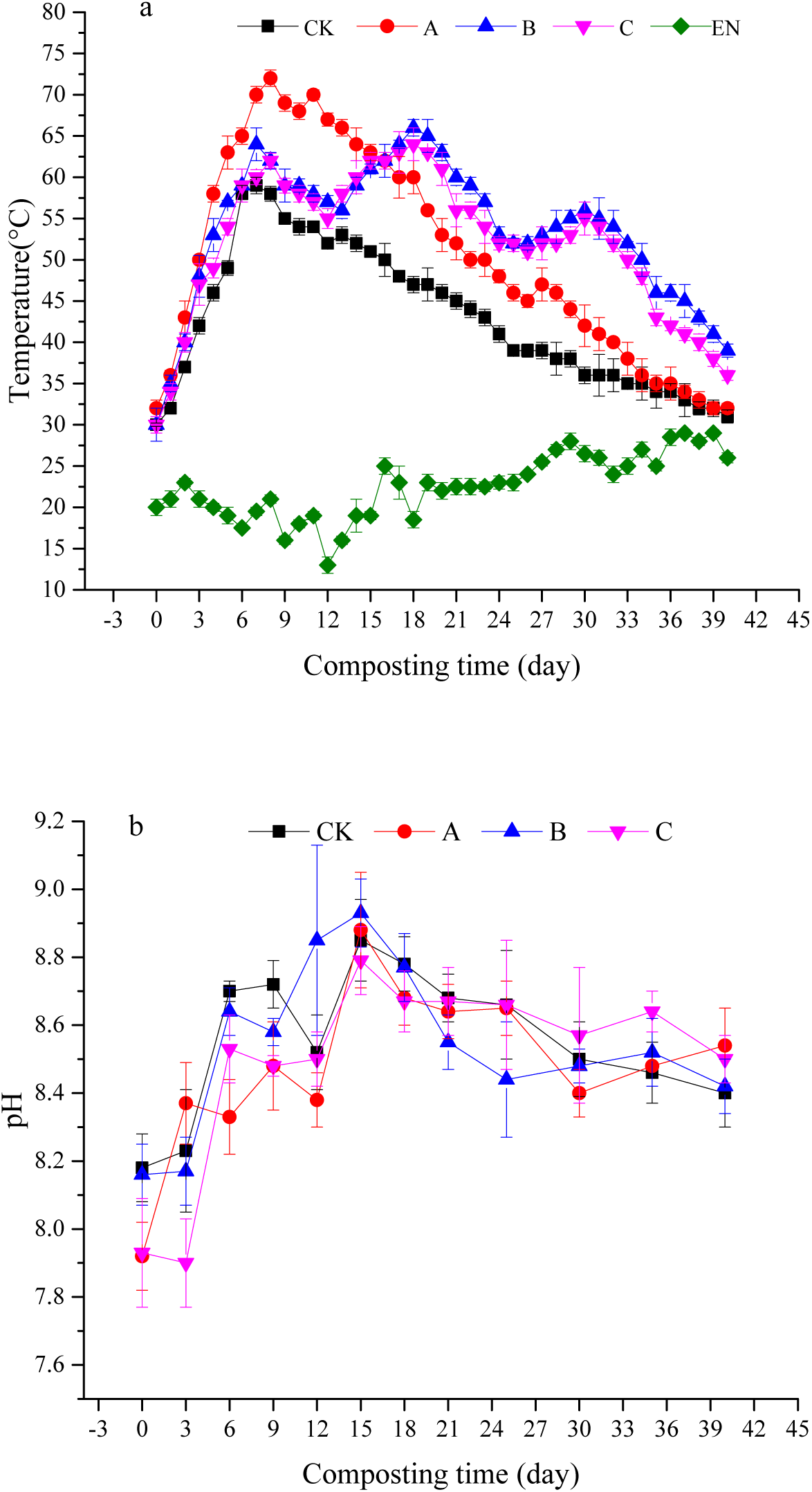
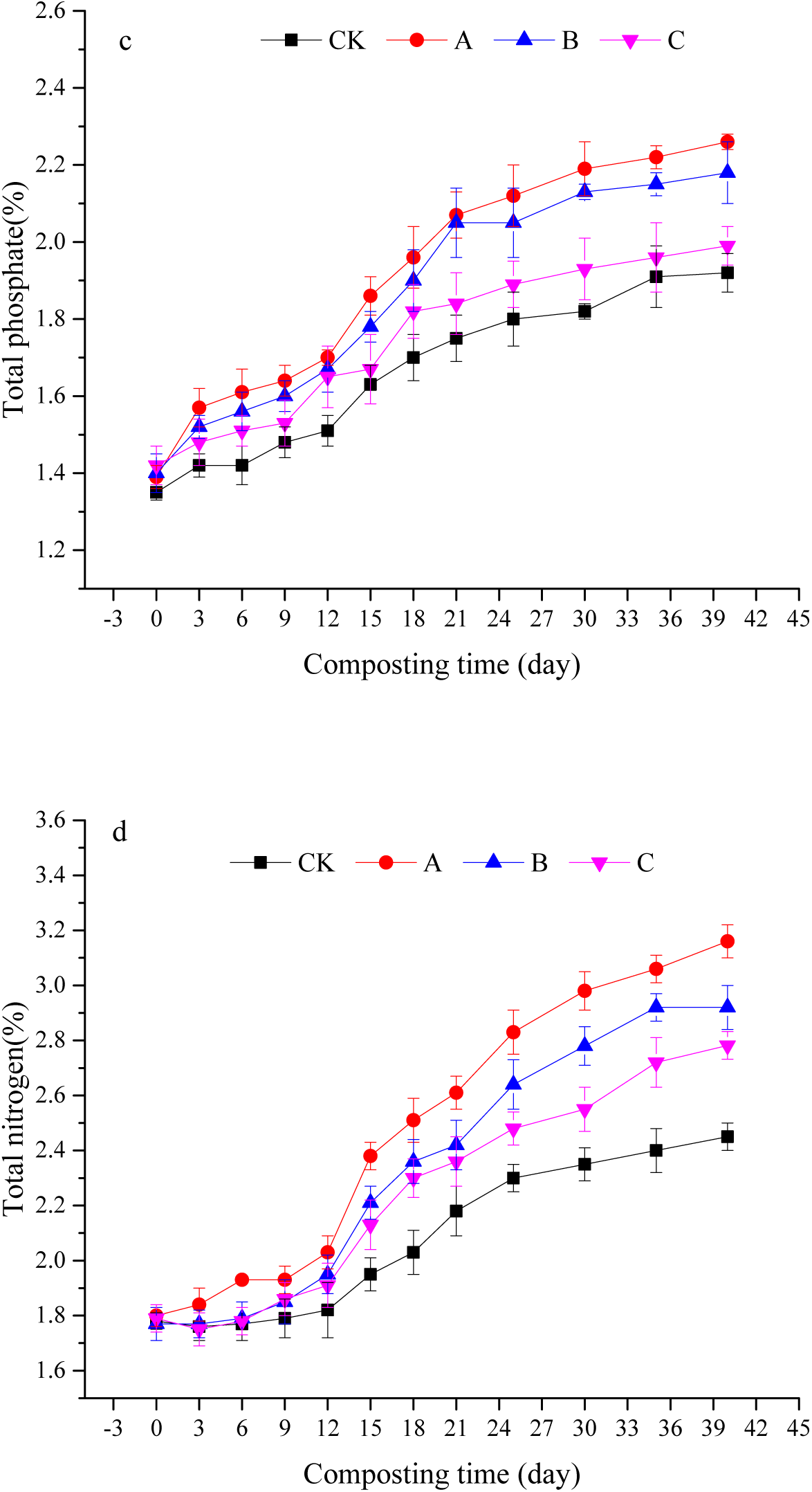
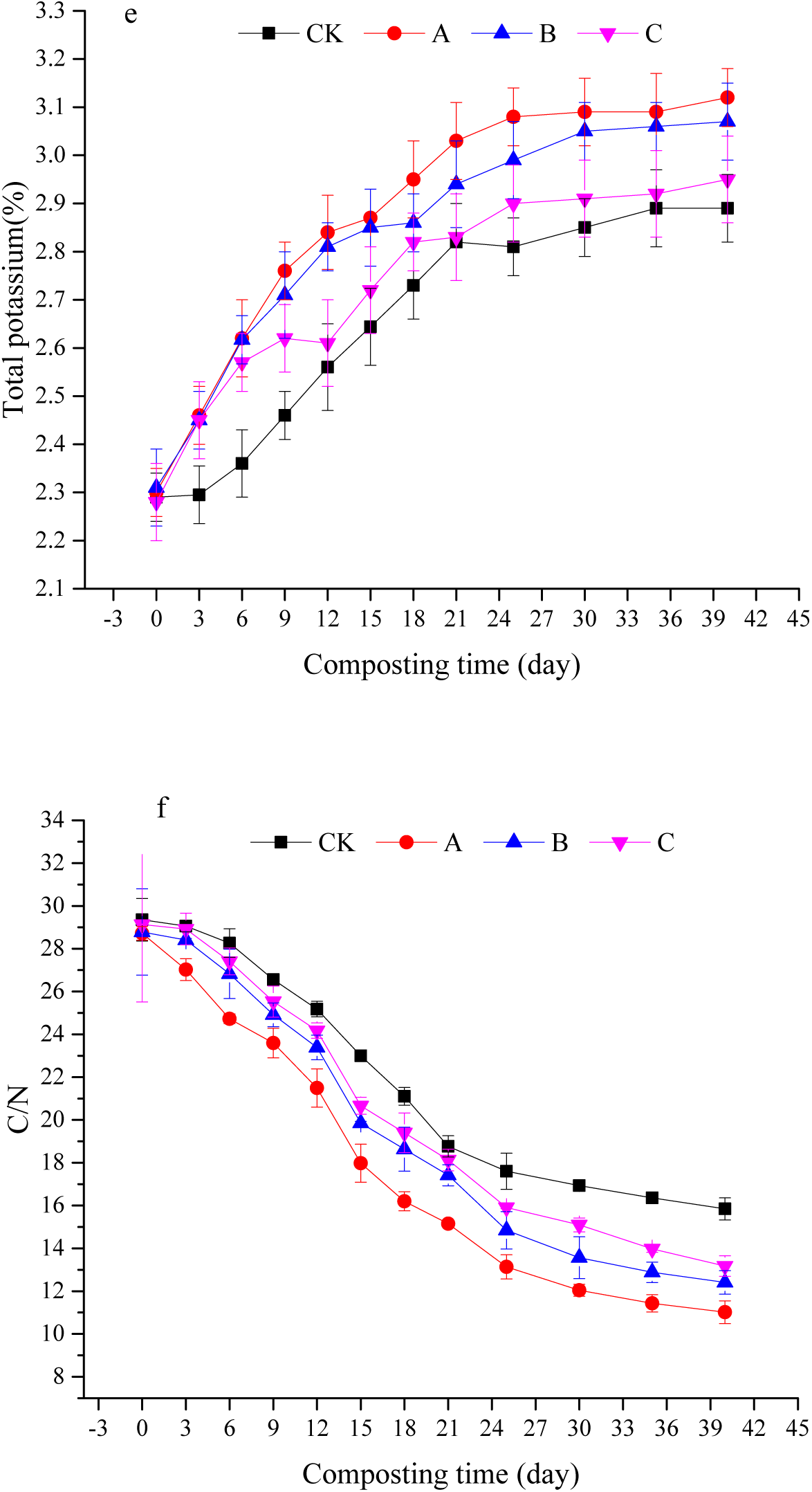
Changes in the physicochemical indices of the four treatment groups during the composting process: a) temperature, b) pH, c) total phosphate, d) total nitrogen, e) total potassium, and f) carbon/nitrogen ratio. EN: Environment temperature.

#### pH changes in compost

pH affects not only microbial metabolic activities, but also mineral redox reactions. The variations in pH are shown in Fig. 1(b). The four groups exhibited similar trends, with an increase to the peak on day 19 followed by a gradual decline. The initial pH in pile A was within the optimal composting pH range (7.0–8.0).

Total nitrogen, total phosphate, and total potassium

Composting is a complex process involving the fixation and release of nitrogen, phosphorus, and potassium, which directly affects compost quality. As composting progressed, TN, TP, and TK showed gradually decreasing trends (Fig. 1c-1e). There were no clear differences in the trends among the four treatments, although pile A (containing M1) exhibited higher levels at all sampling time points. Compared with the effect of commercial microbial agents, higher increases in TN (18.26%), TP (11.54%), and TK (7.89%) contents were observed in group A at the final composting period, greatly exceeding the other treatments. These results showed that agent M1 was conducive to reducing nutritional element, in particular nitrogen.

#### Carbon/nitrogen ratio during composting

Fig. 1(f) presents the changes in the C/N ratio. As composting progressed, the C/N ratio showed a gradually decreasing trend for the initial three weeks and then continued a relatively gentle decline, ranging from 29.35 to 11.01. There were no clear differences in the trends in the C/N ratio among the four groups, although the pile containing M1 showed a greater decrease in the C/N ratio. At the end of composting, pile A had the lowest C/N ratio (11.01), while CK had the highest (15.84).

#### Germination assay

The SGR, RL, and GI values were markedly higher in the piles amended with M1 or commercially available agents than in the unamended piles (Table 2). The maximum SGR, RL, and GI values were 90.48%, 19.32 mm, and 101.60% in treatment A (with M1), and the minimum values were 81.16%, 16.98 mm, and 80.01% in CK. The SGR, RL, and GI values in the pile amended with a mixture of M1 and commercial microbial agents (pile C) were lower than those of A and B, but higher than CK.

**Table 2.** Root length, seed germination rate, and germination index in the four groups.

|  | Root length (mm) | Seed germination rate (%) | Germination index (%) |
| --- | --- | --- | --- |
| CK | 16.98±1.08b | 81.16±2.16b | 80.01±1.01d |
| A | 19.32±0.80a | 90.48±1.08a | 101.6±1.60a |
| B | 18.55±0.93ab | 89.57±2.10a | 95.57±1.27b |
| C | 17.94±1.04ab | 88.21±1.00a | 91.97±2.50c |

Data is reported as mean ± standard error (n = 3); Means with different letters within a column are significantly different at $P < 0.05$ .

### Dynamics of microbial community composition during composting

The 20 samples in this study contained 923,793 sequences and were sorted into 15,118 OTUs. The coverage rate of all samples was over 99%, indicating that the sequencing results were true representations of the microorganisms in the samples (Table 3). As delineated in Fig. A.1, when OTUs were above 30,000, the rarefaction curves reached a plateau. During early composting, pile C had the highest Chao index (998) and Shannon index (5.09), but in the subsequent time points, pile A had the highest values. It suggested that the diversity of microbial community influenced by the different treatment. Each compost environment contained its own unique microbial structure, exhibiting differences in microbial communities during different composting periods.

**Table 3.**
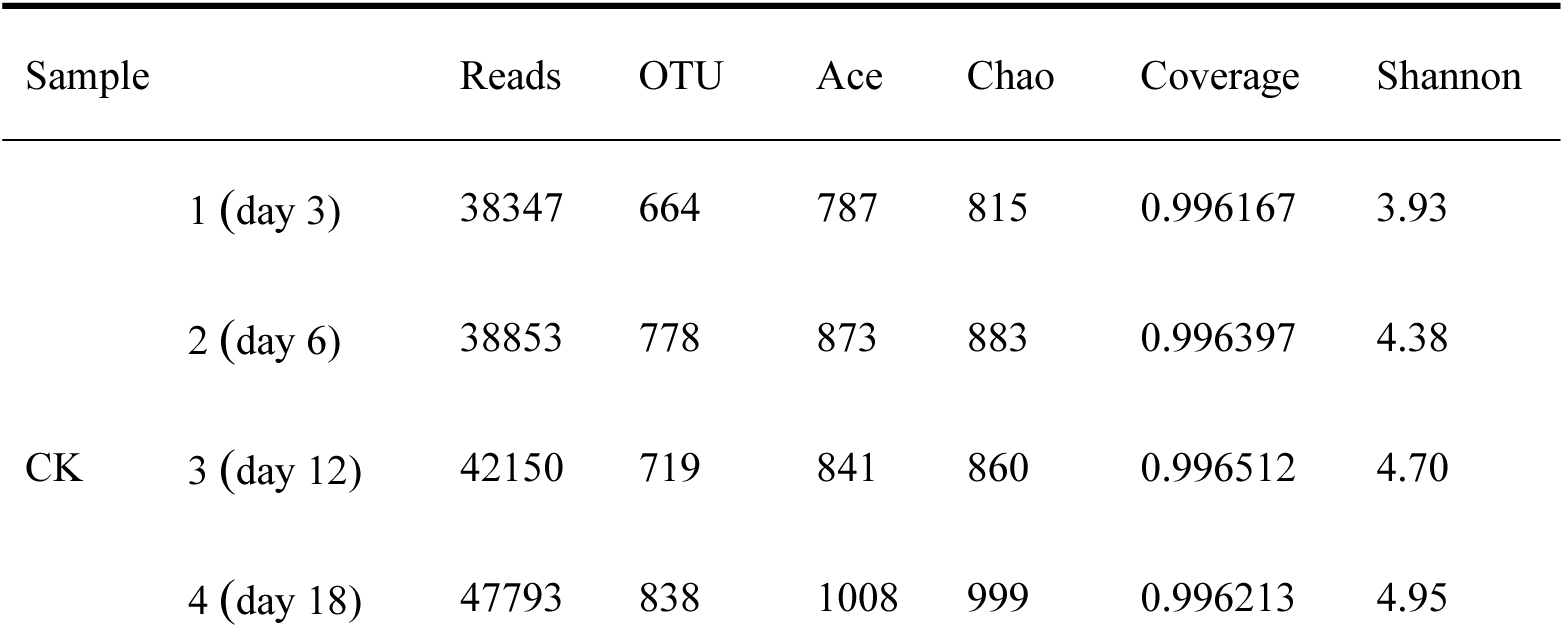
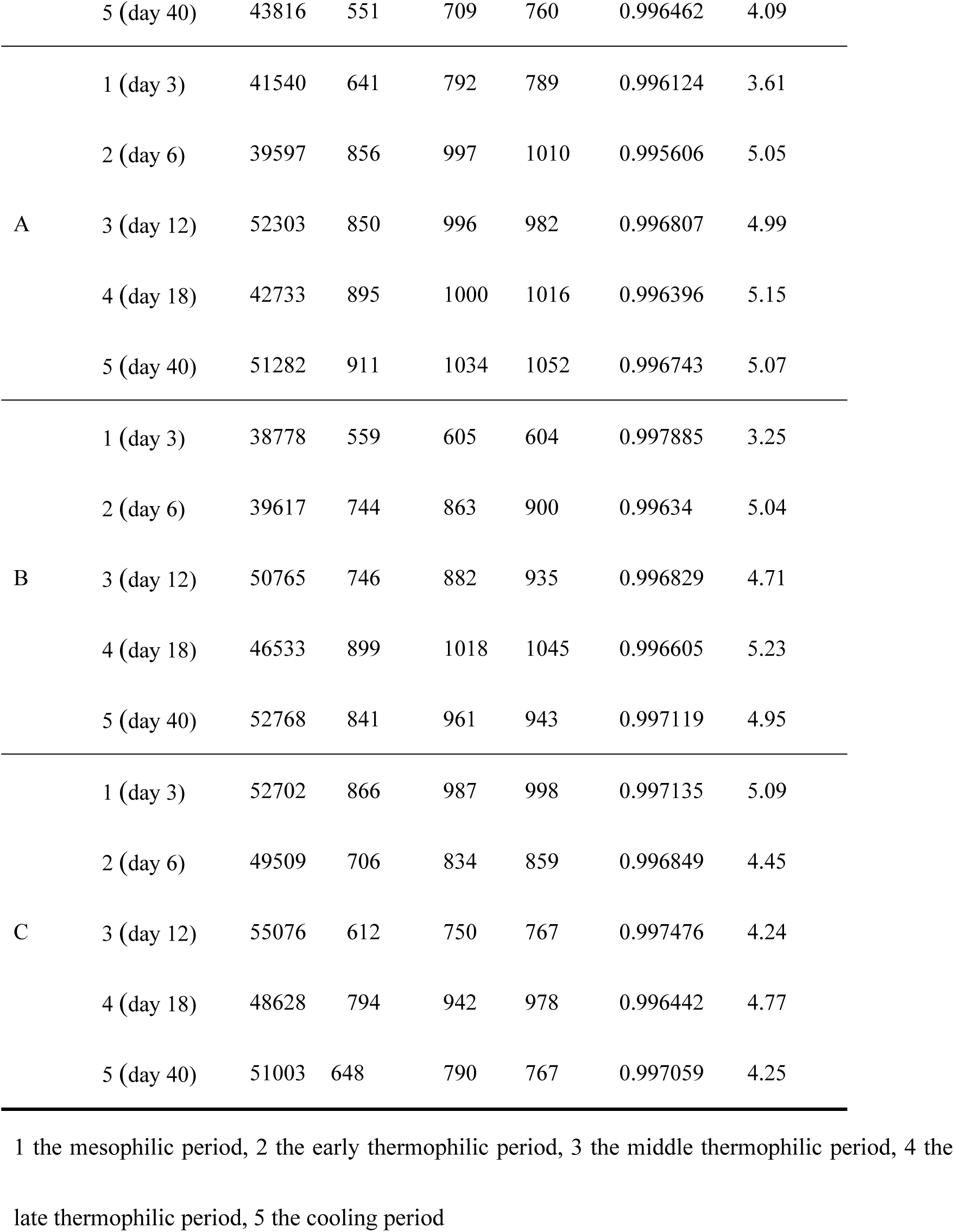
Bacterial richness and diversity indices at 97% similarity level in the four groups.

In order to know the influence of different treatments, non-metric multidimensional scaling (NMDS) was used to examine the variety of microbial community composition. The NMDS plots of the compost samples (Fig. 2) showed that the changes in compost bacterial community compositions were significantly related to NMDS1 and NMDS2, suggesting that treatments could be categorized based on two important factors that affected the microbial community structure. At the end of composting (i.e., late thermophilic and cooling periods), comparing A with B, the compost microbial community composition and structure in A4 and A5 were more similar to those in B4 and B5 than the other treatments, respectively. The dominant phyla in the four groups are shown in Fig. 3. All groups had Proteobacteria, Firmicutes, Actinobacteria, and Bacteroidetes as the dominant phyla. Proteobacteria was the most abundant phylum in the four groups, and the trends of Proteobacteria in groups A and C initially decreased and then increased slightly, whereas B exhibited the opposite trend and CK declined continuously. Firmicutes was the next most abundance phylum, which first increased and then decreased in groups A and C, and increased in CK during the thermophilic period. The abundance of Firmicutes reached a peak during the early thermophilic period in group A (31% of total bacteria) during the mesophilic period in group B (80%), after which the abundance gradually declined. Overall, their abundance remained high during the thermophilic period. Meanwhile, in all four groups Actinobacteria showed the same consistently increasing trend.

**Fig 2.**
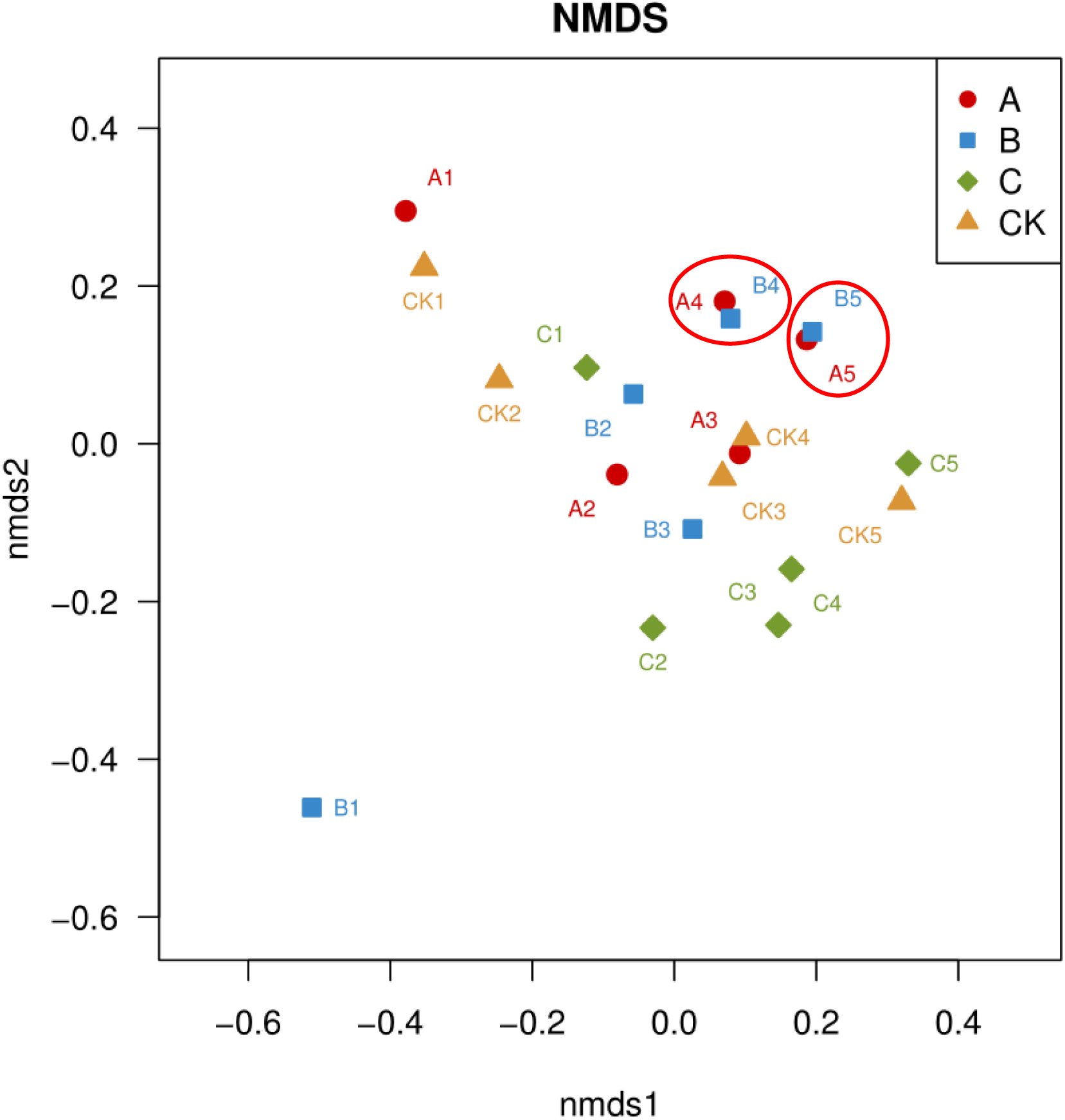
Non-metric multidimensional scaling diagram showing differences in bacterial composition.

**Fig 3.**
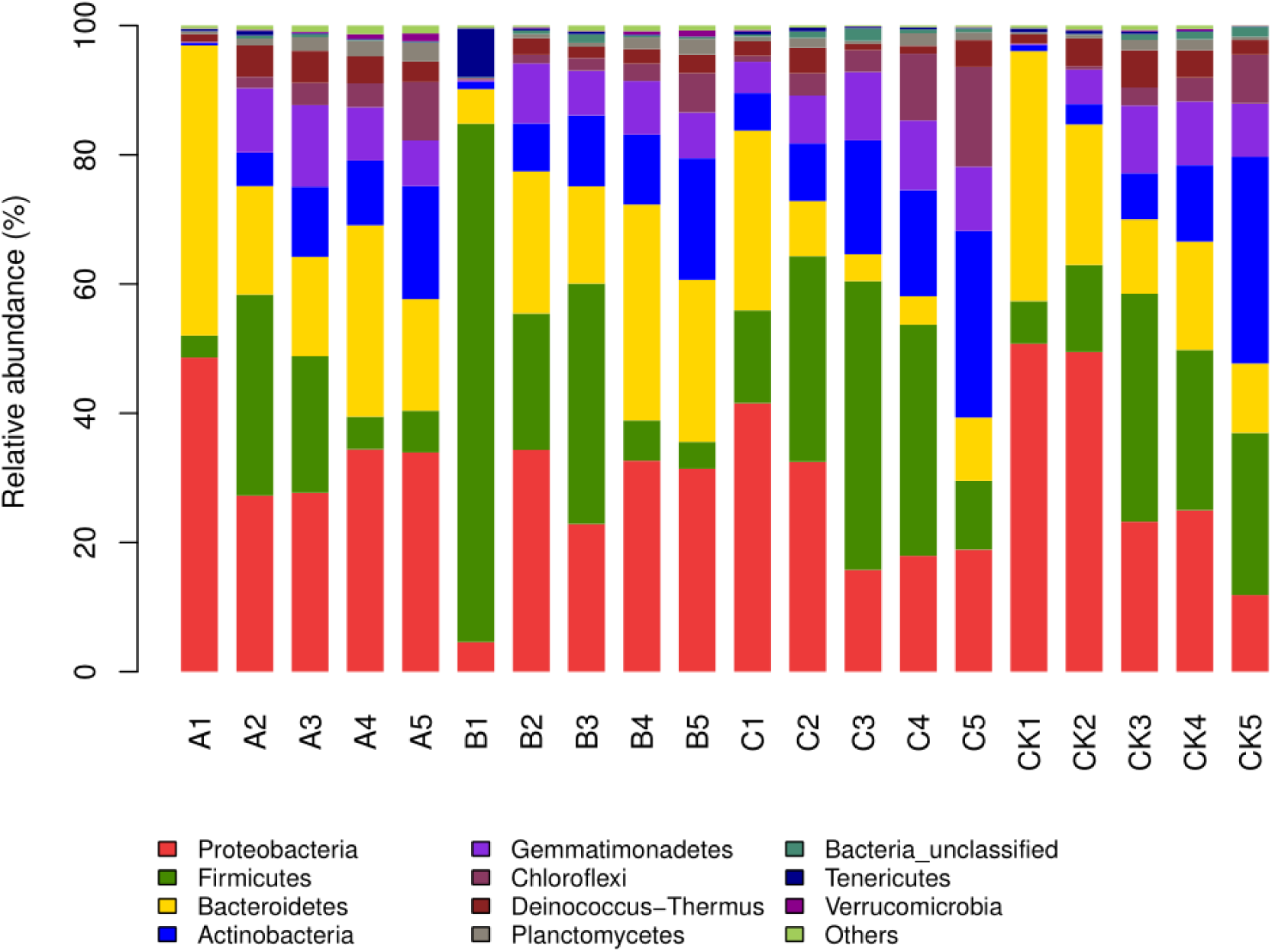
Relative abundances of phyla in each sample.

Fig. 4 shows the total abundances of the five strains in the four groups. In group B, the amount of the five strains decreased during the thermophilic period, in group C, the amount of the five strains in the three stages was relatively low, however, in group A, the abundance of the five strains during the thermophilic period was much greater than in the other treatment groups, ultimately reaching a maximum during the cooling period. The abundance of these strains in all periods was much higher than that in natural compost.

**Fig 4.**
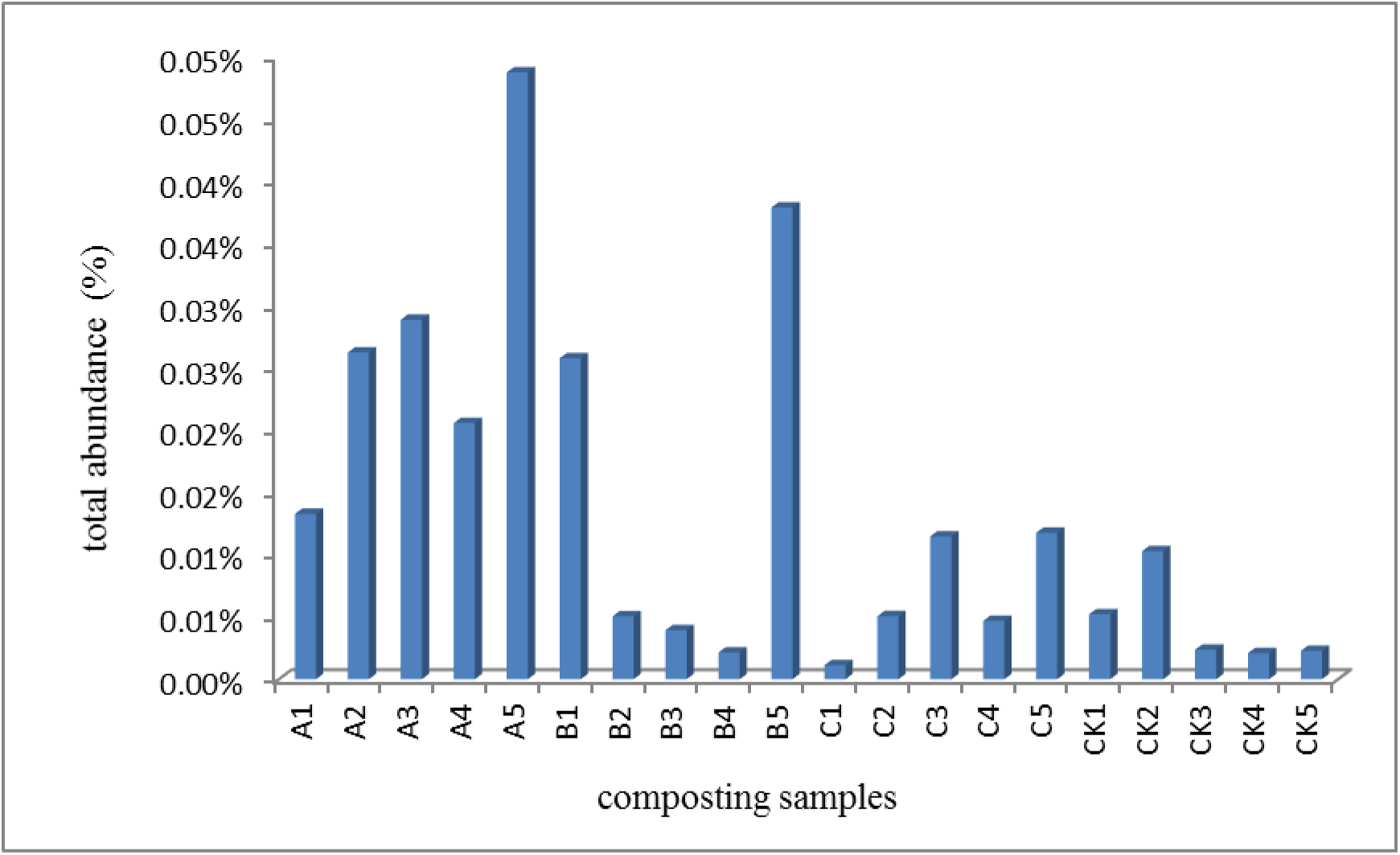
Total abundance of the five *Streptomyces* strains in each composting sample (%).

Redundancy analysis was used to research the physicochemical properties of compost factors on microbial community (Fig. 5). The physicochemical properties were analyzed to determine the key variables responsible for variations in the microbial community at the genus level, considering the top 20 genera. Fig. 5 shows that the first and second components explained 37.77% and 48.70% of the variation in microbial community, respectively. During the cooling period, the microbial community content in groups A, B, C, and CK displayed positive correlations with TP and TK, and the correlation between microbial community in mesophilic period (A, B, C, CK) and C/N ratio was positive.

**Fig 5.**
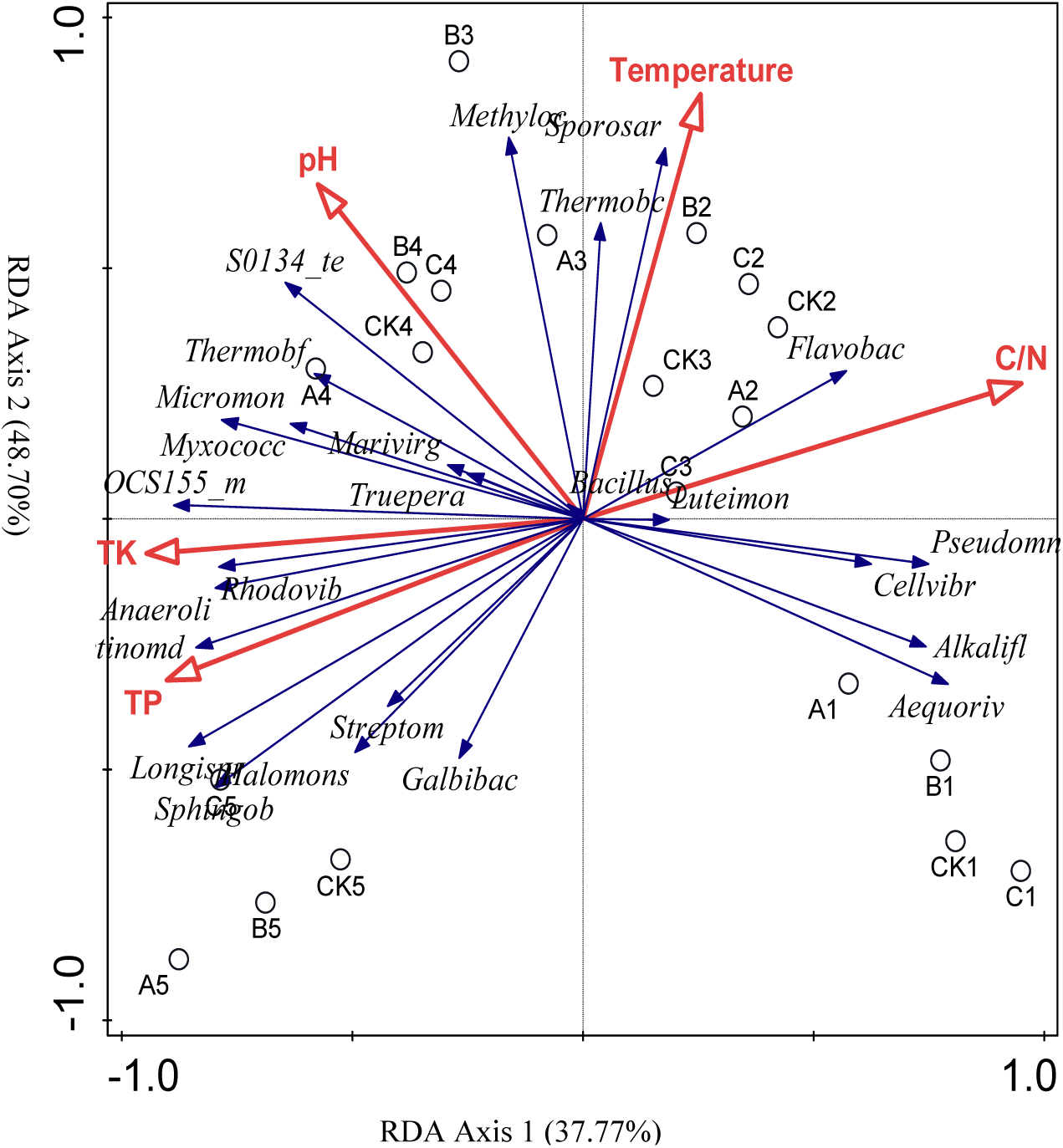
Effects of environmental factors on soil microbial community.

## Discussion

Composting is a biotransformation process under the action of microbe. It is an environmental and economical way for treating garden wastes. While, applying the poor biodegradability of microbial community will lead to low composting efficiency [7,15]. In order to fasten compost maturation process and improve the quality of compost, adding exogenous microorganisms is an effective way. The application of bacteria and fungi were an effectively way which has been proved by many studies [47]. However, few studies have assessed the effect of actinomycetal communities on the composting which peaked during the compost maturation phase. In this study, we applied five high-degradability Actinomycete strains into the composting as additional microbial source for composting of garden waste.

### The effects of microbial agents on chemical and biological indicators

The temperature pattern showed a rapid increase and decreases among all of the treatment. At the beginning of composting, mesophilic microorganisms in the piles decomposed organic matter such as glucose and other carbohydrates and rapidly propagated, which resulted in rising temperatures. However, decreases in readily degradable materials and higher temperatures can reduce microbial metabolism, leading to temperature decreases [20]. The reaction temperature reached 50°C and lasted for over 7 consecutive days, which could adequately reduce pathogens to meet the compost sanitation standard [21]. These results indicated that microbial inoculation accelerated temperature increases, shortened composting cycles, and extended the duration of the high-temperature period. Among the treatments, M1 inoculation exhibited the greatest acceleration in temperature increase.

pH can be used as a measure of the degree of decomposition [21]. The increase may have been due to microbial mineralization of organic matter into simple proteins and production of ammonia, as well as the decomposition of organic acids [22]. The following decrease was probably caused by two mechanisms: (1) an increase in nitrogen loss, (e.g., volatilization of ammonia), and (2) an enhancement in nitrifying bacteria activity to produce organic acids and carbon dioxide [23-24]. The initial pH in pile A was within the 7.0–8.0 pH range, which benefitted microbial activity [25]. And at the end of composting pile A had the lowest pH, which was closer to neutral and suggested a lower impact on soil pH [26].

The contents of TN, TP, and TK increased in all four groups (Fig. 1(c, d, e)), possibly due to the mineralization of organic matter and dry mass reduction [27]. The carbon/nitrogen (C/N) ratio is used extensively to evaluate compost maturity [28-29]. As composting progressed, the C/N ratio showed a gradually decreasing trend for the initial three weeks and then continued a relatively gentle decline, ranging from 29.35 to 11.01. This final value could be considered as within the acceptable degree of maturation according to Wang *et al* [18] This might be due to the increased gaseous loss of carbon as carbon dioxide, while nitrogen remained more tightly bound to organic compounds [30]. Previous studies have shown that higher C/N ratios result in nitrogen volatilization and a reduction in crop growth [31]. Therefore, agent M1 might have induced higher microbial activity and a stronger degradation of complex substrates than the commercial microbial agents and indigenous microorganisms in compost. Compost with a C/N ratio of 12 can be considered as mature [32]. Amendment with agent M1 decreased the C/N ratio of the compost lower than the commercial agent and natural compost, which could have a positive effect on soil fertility [33].

Immature or unstable compost usually has high contents of ammonia, organic acids, or other water-soluble compounds that can inhibit seed germination and plant development. Therefore, safety evaluations of the final compost product are important. The GI is widely used to indicate the phytotoxicity of composting materials. Zuooconi *et al* [34] reported that the GI value can be used as an indication of the phytotoxicity of compost, where the final compost should have a value > 80%. In this study, all final GI values were > 80%. The high SGR, RL, and GI values in piles A (with M1) and B (with commercial microbial agents) may be due to the low generation of short-chain volatile fatty acids by *Streptomyces* spp. Meanwhile, the low SGR, RL, and GI values for pile CK were probably due to the incomplete degradation of toxic acids. High thermophilic temperatures and long thermophilic phases can reduce the phytotoxicity of the compost product [17]. Overall, these results indicated that M1 addition enhanced the biodegradation of toxic materials during composting compared with CK and had similar, or even better, effects compared with the addition of commercial microbial agents. The microbes in the mixture of M1 and commercial microbial agent may have had antagonistic effects, decreasing its safety as a compost product compared to the individual treatments with M1 and commercial microbial agent.

Thus, the application of M1 in composting shortened the composting maturation time and improved the compost quality compared with CK and the commercial microbial inoculum. Inoculated piles had higher decreases in C/N ratio and increases in nutrient contents (TN, TP, and TK), suggesting an increase in compost quality. As such, the five isolated strains have the potential to be applied to large-scale composting.

### The effects of microbial agents on microbial community structure and diversity

The Shannon–Wiener index directly reflects the heterogeneity of a community and species richness [35-36]. The long duration of high temperatures might have deactivated some mesophilic bacterias, while the decrease in the thermophilic period under agent M1 treatment might have alleviated damage to microbes [37]. Comparing treatments A, B, C, and CK, the microbial diversity and abundance of group A were the highest among the four groups. This indicated that adding M1 could increase the dominant bacteria diversity, and decrease the competition between the inoculants and indigenous bacteria. This further illustrates that thermotolerant microorganisms are the biological basis of compost and that exogenous agents are necessary to improve the maturation and efficiency of inoculation composting [38]. At late thermophilic and cooling periods, the compost microbial community composition and structure between treatment A and B were similar. This suggested that the dominant microbial communities at the OTU level in groups A and B at the end of composting were similar to those found in the commercial microbial inoculum, which are used to accelerate the composting process, indicating that M1 altered the overall microbial community structure and maintained the microbial richness and diversity of the compost. In group C, the microbial communities at the OTU level differed from the communities of A and B during the mesophilic and cooling periods, indicating possible antagonism or competition between the two agents [39].

The relative abundances of the dominant microbes at the phylum level can comprehensively reflect the community structure and diversity of microorganisms in different habitats [40-41]. The dominant phyla in the four groups are Proteobacteria, Firmicutes, Actinobacteria, and Bacteroidetes, consistent with previous research [18,42,43]. Proteobacteria have the primary function of the degradation of soluble sugar and fixed nitrogen [44-46]. The trends of Proteobacteria in groups A and C initially decreased and then increased, suggesting agent M1 stimulated the growth of Proteobacteria during the cooling period, potentially improving compost maturation. Firmicutes was first increased and then decreased in groups A and C, and increased in CK during the thermophilic period. Firmicutes are the main bacteria responsible for hydrolyzing polysaccharides, such as cellulose, which is critical for the subsequent composting steps, which may contribute to their survival strategy in hot environments [47-48]. Agent M1 increased the abundance of Firmicutes during the initial thermophilic period, promoting the decomposition of organic matter. In all four groups Actinobacteria showed increasing trend, consistent with the results of Wang *et al* [10], Cheng *et al* [15], and Zhao *et al* [13] During the composting process, the steady rise in Actinobacteria highlighted their degradation functions during the thermophilic and cooling phases, possibly due to their thermostability and the direct correlation with lignocellulose and humus contents [49-50]. Zimmermann *et al* [51] found that a diverse range of mesophilic and thermophilic actinomycetes in Actinobacteria could degrade lignocelluloses. The trends in Bacteroidetes abundance in group A were similar to those in group CK, which only significantly increased after the middle thermophilic period. Bacteroidetes mainly degrade the substrates remaining after pile temperatures drop [52] and agent M1 appeared to enhance this effect compared to natural composting.

During the mesophilic composting phase, mesophiles such as Proteobacteria and Bacteroidetes broke down organic matter rapidly for energy production and growth, generating heat and inducing changes in the C/N ratio [53]. When entering the thermophilic period, high temperatures inhibited mesophile growth, and thermophiles such as Firmicutes and Actinobacteria had major roles in decomposing cellulose and producing humus. At the same time, NH_4_-N was converted into NH_3_, which could then be volatilized, increasing the pH. Thermotolerant microbes could also increase NO_3_-N concentrations to maintain total nitrogen levels [54]. Once the temperature began to drop, some mesophiles became active again and further matured the piles [33]. Physicochemical characteristics (e.g., temperature, pH, and element contents) of the piles undergo dynamic changes according to the succession of microbial communities [57]. The physicochemical properties were analyzed to determine the key variables responsible for variations in the microbial community at the genus level, considering the top 20 genera. Redundancy analysis showed that different factors limited the members of the microbial community in different stages at the genus level, and that the ecological habitat had a significant effect on bacterial abundance. Interestingly, *Streptomyces*, as found in M1, was negatively correlated with C/N and positively correlated with TP and TK. For example, the abundance of the five strains reached a maximum, along with TN and TP, during the cooling period, when the C/N ratio was low. These results directly showed that adding agent M1 significantly improved garden waste composting and altered the physicochemical parameters to induce shifts in actinomycetal communities.

The results of this study showed that the combination of five strains (M1) improved the degradation of organic matter and enhanced the composting speed. The total abundances of the five strains in group B decreased during the thermophilic period, probably due to the death of some genera by high temperatures. Meanwhile, in group C, which combined M1 with the commercial microbial inoculum, the amount of the five strains in the three stages was relatively low, possibly due to resource competition between M1 and the commercial microbial inoculum. We speculated that M1 activity began at the beginning of the mesophilic period, and degraded cellulose during the thermophilic period, which influenced the microbial community structure, and subsequently the characteristics, of the pile [55-56].

## Conclusions

From a macroscopic perspective, compared with the commercial microbial inoculum (treatment B), agent M1 not only resulted in more complete organic matter degradation and enhanced composting speed, but also increased the fertility of the resulting compost. From a microscopic perspective, the structure of the microbial communities in the groups with M1 and the commercial microbial inoculum were similar, indicating that M1 adjusted the overall microbial community structure and maintained the microbial richness and diversity in the compost. Therefore, application of agent M1 to compost can accelerate the composting process and shorten the composting period.

## Conflict of interest

The authors declare that they have no conflict of interest.

## Acknowledgements

We thank at least two professional editors from TEXTCHECK (http://www.textcheck.com/certificate/GBPkw5), for their English improvement to the paper. This work was financially supported by National Undergraduate Training Programs for Innovation and Entrepreneurship (2017100220206); the Beijing Municipal Science and Technology Project (Z151100002115006); National Undergraduate Innovation and Enterpreneurship Training Program (No.201710022026).

---

Fig S1. Rarefaction curves based on high-throughput sequencing of bacterial communities from four groups at a cutoff level of 97%.

